# The CRY1 tail controls circadian timing by regulating its association with CLOCK:BMAL1

**DOI:** 10.1101/758714

**Authors:** Gian Carlo G. Parico, Ivette Perez, Jennifer L. Fribourgh, Britney N. Hernandez, Hsiau-Wei Lee, Carrie L. Partch

## Abstract

Circadian rhythms are generated by a transcription-translation feedback loop that establishes cell-autonomous biological timing of ~24-hours. A prevalent human variation in the core clock gene cryptochrome 1, Cry1Δ11, lengthens circadian period to cause Delayed Sleep Phase Disorder (DSPD). CRY1 has a 55 kDa photolyase homology region (PHR) followed by a ~100 residue tail that is intrinsically disordered; the Δ11 variant lacks a short segment encoded by Exon 11 within its tail. We show here that the disordered tail of CRY1 interacts directly with its PHR, and that Exon 11 is necessary and sufficient to disrupt the interaction between CRY1 and CLOCK, a subunit of the primary circadian transcription factor. Competition between PER2 and the tail for the CRY1 PHR suggests a regulatory role for the tail in the early morning, when CRY1 binds to CLOCK:BMAL1 on DNA independently of PER2. Discovery of this autoregulatory role for mammalian CRY1 highlights functional conservation with plant and insect cryptochromes, which also utilize PHR-tail interactions to reversibly control their activity.

**One Sentence Summary:** The disordered tail of the CRY1 protein regulates interactions between CRY1 and other core circadian rhythm proteins.

## Main Text

Circadian rhythms coordinate behavior and physiology with the 24-hour solar day. At the molecular level, over 40% of the genome is temporally regulated in a circadian manner (1). At the core of this rhythmic gene expression, the heterodimeric transcription factor CLOCK:BMAL1 (brain and muscle Arnt-like protein 1) promotes transcription of its own repressors, period (PER1/2) and cryptochrome (CRY1/2) (2). We previously showed that changing the affinity of these regulators for one another can alter circadian timing by shortening or lengthening the duration of repression in the feedback loop (3, 4). As a consequence, variations within these core circadian genes can lead to sleep disorders such as DSPD, which is characterized by a longer than normal circadian period (>24.2h) and later than normal sleep/wake times (5, 6).

One form of familial DSPD arises from a variant of CRY1 (CRY1Δ11) that is as common as 1 in 75 members of certain populations (7). CRY1Δ11 disrupts a splice site leading to the in-frame deletion of exon 11, thus removing 24 residues within the C-terminal tail of CRY1. While the tail is not necessary for CRY1 to interact with CLOCK:BMAL1 and generate circadian rhythms, its truncation or modification affects the periodicity and amplitude of rhythms (7–10). By contrast, the CRY1 PHR is required for circadian rhythms and binds directly to the CRY-binding domain (CBD) of PER1/2 as well as both CLOCK and BMAL1 subunits (3, 11, 12). We showed earlier that the tighter CRY1 binds to CLOCK:BMAL1, the longer the circadian period becomes (3, 13). This suggested that the CRY1Δ11 variant, which enhances co-immunoprecipitation with CLOCK:BMAL1 and extends the circadian period (7), may alleviate an autoinhibitory interaction between the CRY1 tail and PHR to increase its affinity for CLOCK:BMAL1. CRY1Δ11 therefore appears to mimic mutants in Drosophila cryptochrome that are constitutively active due to truncation of their C-terminal tail (14, 15), in contrast to full-length cryptochromes that are autoinhibited through a tail/PHR interaction (16, 17).

To determine whether the CRY1 tail interacts with the CRY1 PHR, we used fluorescence polarization (FP) assays to explore binding of a fluorescently labeled human CRY1 tail to the PHR in trans (Figure 1A). The resulting binding curve is consistent with single-site protein-ligand binding and fits to an equilibrium dissociation constant (K_D_) of about 5 µM (Figure 1B-C). The Δ11 version of the hCRY1 tail interacts with about four-fold lower affinity, demonstrating that exon 11 plays an important role in binding the PHR (Figure 1B,C). However, a peptide encoding the isolated exon 11 also binds with lower affinity, suggesting that additional sites on the CRY1 tail contribute to PHR binding. Using an exon-based truncation analysis, we determined that exons 10 and 11 (residues 496-554) comprise the minimal binding region of the hCRY1 tail for the PHR (Figure 1B,C). The mouse CRY1 tail, which contains a short repeat insertion in exon 10, binds to its PHR with a similar affinity to the human tail (Supplementary Figure 1). Finally, our data suggest that exon 12 does not enhance affinity for the PHR (Figure 1B,C). This was surprising since post-translational modification of this region regulates intracellular levels of CRY1 to play a role in circadian timekeeping (9, 18). Phosphorylation of S588 in mouse CRY1 (S568 in humans) after DNA damage increases its stability, as does the corresponding phosphomimetic mutation S588D (9, 18). Consistent with our observation that exon 12 does not contribute to PHR binding, we found no change in affinity of the S588D CRY1 tail for the PHR (Supplementary Figure 1).

**Fig. 1.:**
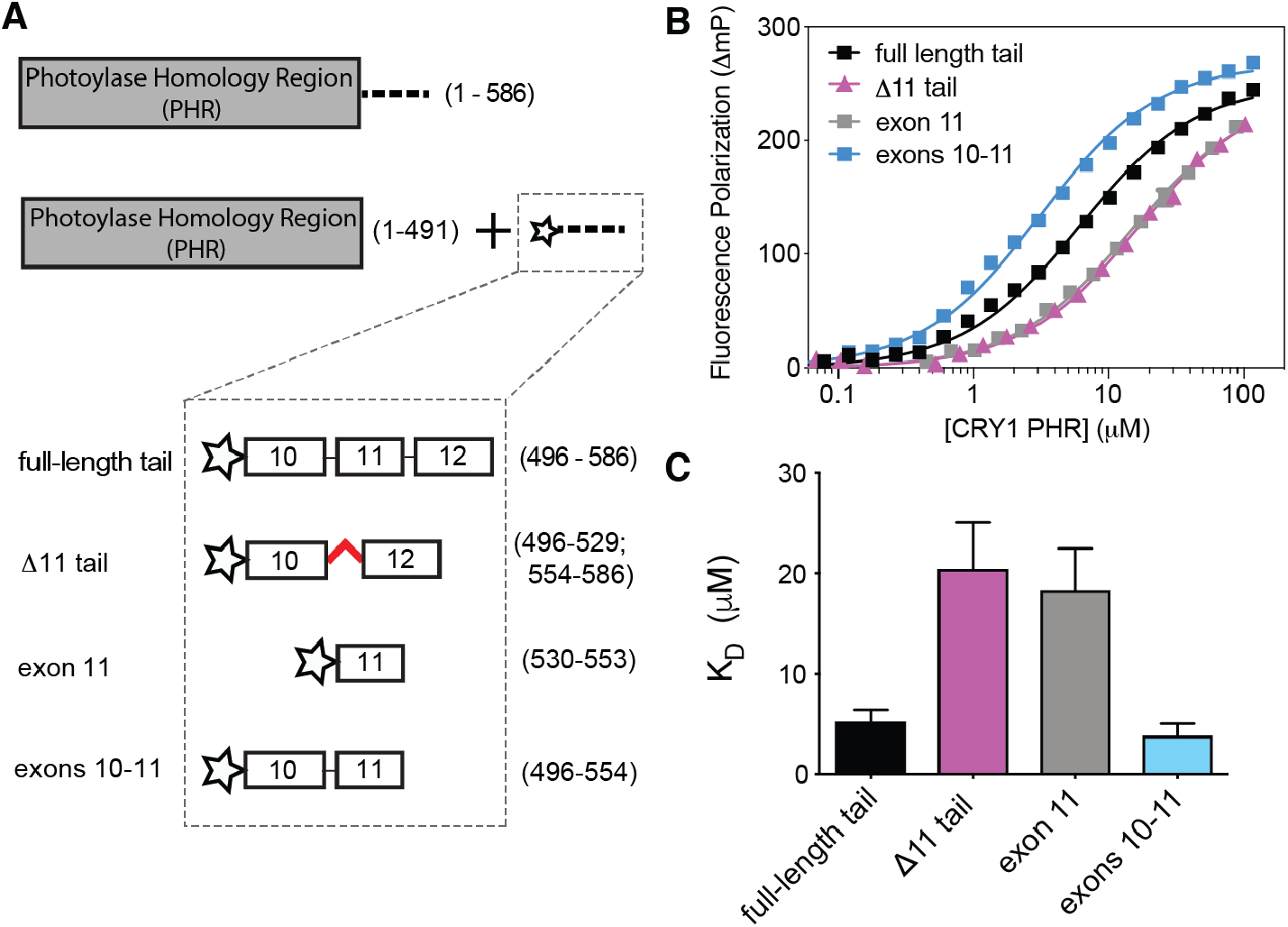
Exons 10 and 11 in the CRY1 tail are required for binding to its PHR domain. (A), Domain architecture of full-length human CRY1 and the constructs used in FP experiments, comprising the PHR (residues 1-491, gray box) and tail (residues 492-586, dashed line) with various truncations; the N-terminal fluorescein (FAM) label is depicted as a star. (B), FP binding curves of fluorescently-labeled human tail constructs to the CRY1 PHR. Plot shows the mean representative binding curves of duplicate samples ± sd (of n = 3 independent assays). Curve represents fit to one-site binding (Prism). (C), Affinities of FAM-tail constructs for the PHR derived from FP binding assays (n = 3 independent assays ± sd).

To further map the PHR-binding epitopes on the tail, we used nuclear magnetic resonance (NMR) spectroscopy to probe the CRY1 tail with atomic resolution. Backbone-based ^1^H-^15^N HSQC experiments probe the chemical environment of all non-proline residues as individual peaks; however, these spectra exhibit severe peak overlap in the case of long intrinsically disordered proteins (IDPs) such as the CRY1 tail (Figure 2A) (19, 20). To circumvent this problem, we turned to ^13^C carbon-detected methods such as the ^13^C-^15^N CON to increase peak dispersion and therefore, our resolution, on this IDP (21). The CON spectrum of the CRY1 tail exhibited excellent peak dispersion that allowed us to visualize and assign 81% of the backbone chemical shifts (Supplementary Figure 2).

**Fig. 2.:**
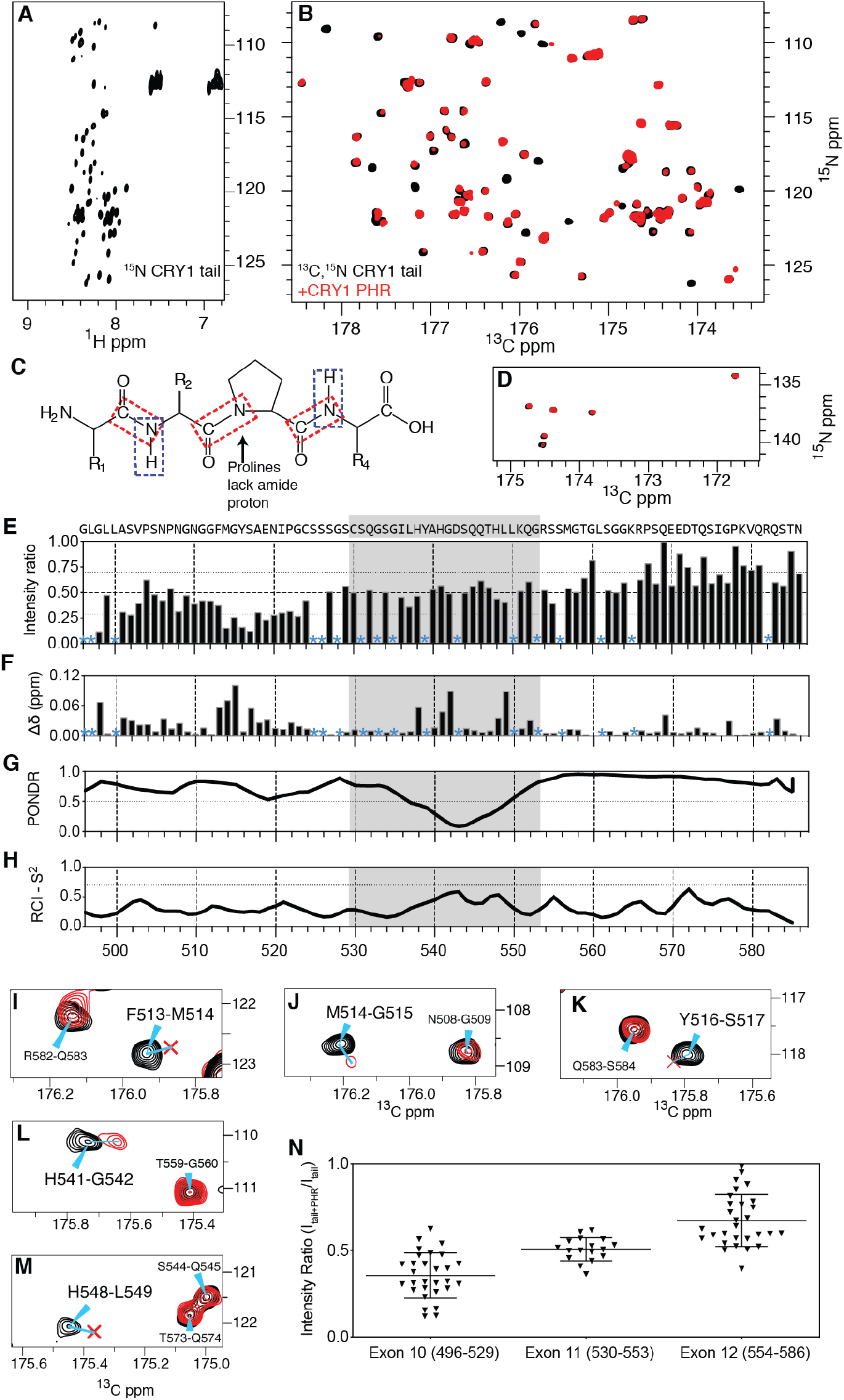
NMR spectroscopy maps the PHR binding site on the disordered CRY1 tail. (A), ^1^H-^15^N HSQC spectrum of the isolated ^15^N-labeled CRY1 tail at 120 µM. (B), ^13^C-^15^N CON spectrum of the ^13^C, ^15^N-labeled CRY1 tail at 120 µM alone (black) or in the presence of 120 µM CRY1 PHR (red). (C), Schematic of proline-containing peptide backbone, highlighting the correlations visualized by ^1^H-^15^N HSQC (blue box) or ^13^C-^15^N CON (red box). (D), Proline-specific region of the ^13^C-^15^N CON spectrum of the ^13^C, ^15^N-labeled CRY1 tail ± CRY1 PHR (as in panel B). (E-F), Chemical shift perturbation (CSP, E) and relative intensities (F) in the ^13^C, ^15^N-labeled CRY1 tail upon addition of equimolar CRY1 PHR. Asterisk, unassigned or overlapping peaks excluded from the analysis. Exon 11 is represented by a gray box. In H, dashed horizontal line represents the mean intensity ratio (I_tail+PHR_/I_tail alone_), while the dotted lines represent sem. Peaks are referred to by the number of the nitrogen of the ^13^C-^15^N peptide bond. (G), PONDR prediction of a disorder in the CRY1 tail with an ordered minimum in Exon 11. (H), CSI prediction of the lack of secondary structure from NMR data (S^2^ < 0.7). (I-M), Zoom in on ^13^C-^15^N CON spectra peaks (residues F513, M514, G515, Y516, H541, H548 respectively) that are severely broadened or perturbed upon addition of CRY1 PHR. (N), Intensity ratios grouped by Exon (mean ratio per exon ± sd).

To probe the role of exon 11 in the CRY1 tail/PHR interaction, we added CRY1 PHR to the ^13^C,^15^N-labeled CRY1 tail and monitored chemical shift perturbations. We were limited to some degree by the solubility and stability of the CRY1 PHR under our NMR conditions, but under equimolar concentrations of tail and PHR, we identified a number of peaks that exhibited PHR-dependent changes indicative of binding (Figure 2B). Another advantage of carbon-detected methods is their ability to visualize proline chemical shifts, often enriched in IDPs, that cannot be detected using standard ^1^H-^15^N HSQC spectra (Figures 2C,D). For the most part, we observed broadening at select peaks within the ^13^C,^15^N-labeled CRY1 tail upon binding to the PHR, likely due to the large size of the PHR (~55 kDa) (Figure 2B,D-E). However, some clusters of peaks in exon 10 and 11 demonstrated changes in chemical shift (Figure 2F). Of note is a hydrophobic patch in exon 10 that is composed of the motif Phe-Met-Gly-Tyr and several histidines in exon 11 (Figures 2I–M and Supplementary Figure 2). These same peaks are also broadened relative to the rest of the tail, suggesting that these residues are hotspots for interacting with the PHR (Figure 2E). An exon-based analysis of peak broadening demonstrates that it occurs to the highest extent in exons 10-11, and to a lesser degree in exon 12 (Figure 2N), corroborating our FP data that identified exons 10-11 as the minimal binding region on the tail for interacting with the PHR.

We also utilized the NMR data to probe structural characteristics of the isolated CRY1 tail. The narrow proton chemical shift dispersion and overlap of peaks in the ^1^H-^15^N HSQC spectrum is certainly representative of an IDP (Figure 2A), in agreement with circular dichroism data of the CRY1 tail published earlier (16). Although the computational tool PONDR predicted that the CRY1 tail is highly disordered, it suggested that there may be some propensity for structure within an ordered ‘minimum’ in exon 11 (Figure 2G) (22); these local, ordered minima have been implicated as linear binding epitopes within IDPs (23). However, the chemical shift index (CSI) derived from NMR data predicts that the entire CRY1 tail lacks order independent of the PHR (Figure 2H) (24). Further studies could identify if any regions in the tail, such as exon 11, might adopt secondary structure upon binding to the PHR.

Given that exon 11 plays an important role in the interaction between the CRY1 tail and its PHR, we next wanted to determine if the CRY1 tail regulates binding between the CRY1 PHR and CLOCK:BMAL1. We previously identified two distinct interfaces for interaction with the CLOCK:BMAL1 heterodimer on the CRY1 PHR: the coiled-coil (CC) helix of CRY1 binds to the transactivation domain (TAD) of BMAL1 (3, 16), while the secondary pocket of CRY1 binds directly to CLOCK PAS-B (Figure 3A) (12). We established FP-based binding assays for the PHR using fluorescently labeled BMAL1 TAD (3) or CLOCK PAS-B (Supplementary Figure 3), allowing us to form complexes of CRY1 PHR with either target and determine the ability of exogenous tail to compete for interaction on the PHR in trans. Titrating CRY1 tail into a CRY1 PHR:BMAL1 TAD complex did not result in any changes in fluorescence polarization, demonstrating that the CRY1 tail likely does not play a role in regulating the interaction between the CRY1 PHR and the BMAL1 TAD (Figure 3B). However, titrating full-length CRY1 tail to a CRY1 PHR:CLOCK PAS-B complex caused a dose-dependent decrease in FP signal to levels observed with the isolated CLOCK PAS-B, demonstrating that the tail displaced CLOCK PAS-B from the CRY1 PHR (Figure 3C). Furthermore, we found that a peptide comprising the isolated exon 11 also displaced CLOCK PAS-B from the CRY1 PHR, while the Δ11 tail had no effect on the CRY1 PHR:CLOCK PAS-B complex (Figure 3C). Therefore, exon 11 is necessary and sufficient to regulate the interaction between CLOCK PAS-B and the CRY1 PHR.

**Fig. 3.:**
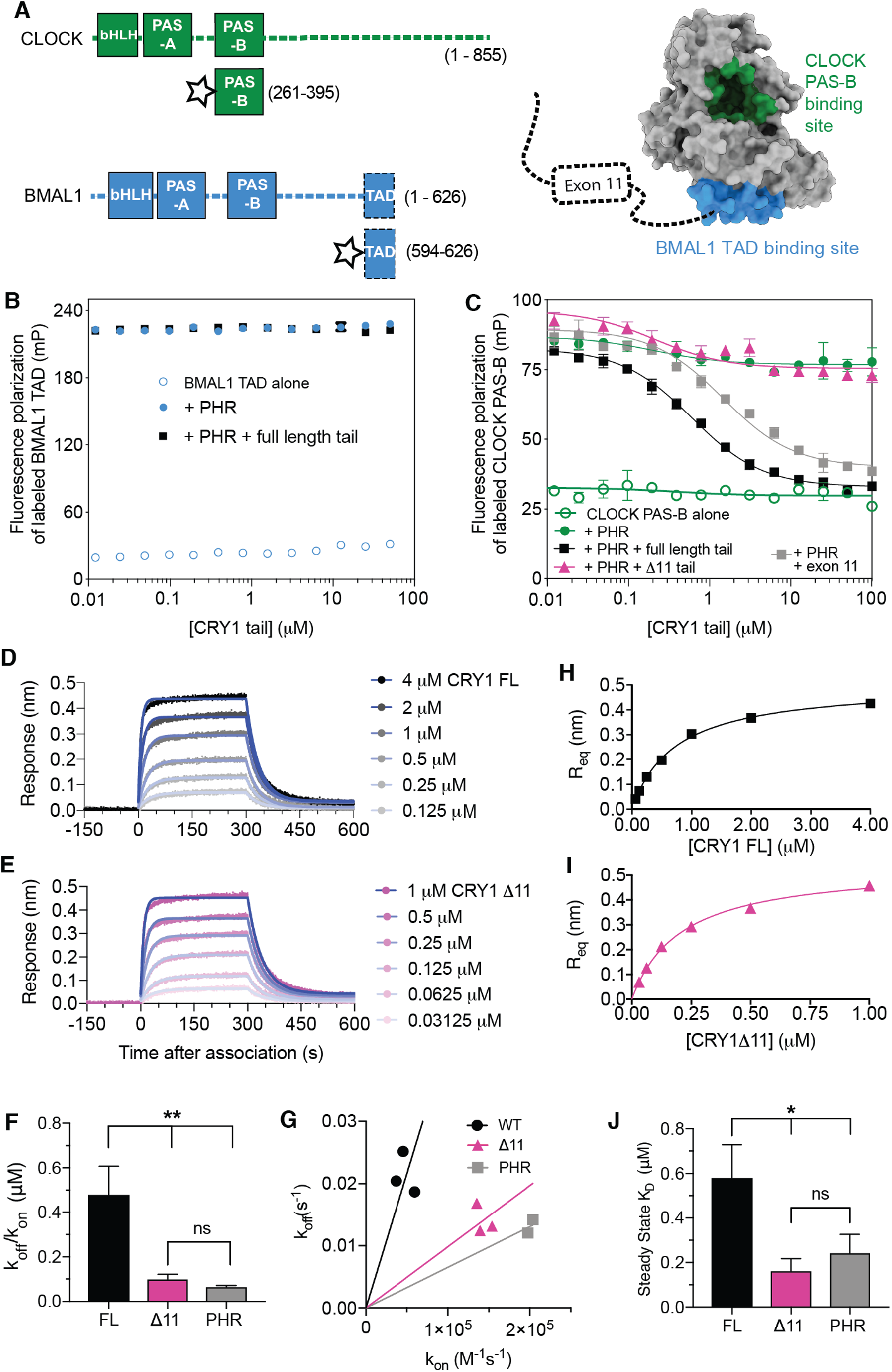
Exon 11 competes with CLOCK PAS-B to reduce CRY1 affinity for CLOCK:BMAL1. (A), Domain architecture of CLOCK and BMAL1. Starred constructs represent those used for FP assays. Right, CLOCK (green) and BMAL1 (blue) binding sites mapped onto the CRY1 PHR (PDB: 5T5X). (B), FP-based equilibrium competition assay with 4 µM CRY1 PHR bound to 20 nM fluorescently-labeled BMAL1 TAD in the absence (filled blue) or presence of CRY1 tail (black). Data represent mean ± sd from one representative assay (of n = 3). (C), FP competition assay with 4 µM CRY1 bound to 20 nM fluorescently-labeled CLOCK PAS-B (filled green) in the presence of CRY1 tail (black), exon 11 (gray), or the Δ11 tail (magenta). Data represent mean ± sd from one representative assay (of n = 3). For assays with displacement, curves represent fit to one-site competitive binding model (Prism). (D-E), BLI sensorgram for biotinylated CLOCK:BMAL1 PAS-AB titrated with full-length CRY1 (E, gray to black) or CRY1Δ11 (pink to magenta). Model fit represented by thin blue line. (F), Fits for K_D_ from kinetic analysis (k_off_/k_on_) of BLI data. (G), Plot of k_on_ vs. k_off_ for CRY1 constructs analyzed by BLI. (H-I), Steady-state analysis of BLI response versus CRY1 full-length (black) or CRY1Δ11 (magenta) concentration (from data in panel D and E, respectively). (J), Fits for K_D_ from steady-state analysis of BLI data.

One prediction of our model is that both CRY1Δ11 and the tail-less CRY1 PHR should have higher affinity for the CLOCK:BMAL1 PAS domain core compared to full-length CRY1. We used Biolayer Interferometry (BLI) to quantitatively assess CRY1 binding to immobilized CLOCK:BMAL1 PAS-AB heterodimer. Importantly, even though other PAS domains are present in the CLOCK:BMAL1 PAS-AB heterodimer, our previous studies showed that association with CRY1 absolutely depends on CLOCK PAS-B docking into the secondary pocket of CRY1 (Supplementary Figure 3) (12). We tracked the binding kinetics (association and dissociation) of CRY1 over time (Figure 3D, E) to confirm that both CRY1Δ11 and the isolated PHR have a higher affinity for the CLOCK:BMAL1 PAS domain core than full-length CRY1 (Figure 3F). Analysis of the kinetic binding data suggested that the decrease in affinity of full-length CRY1 for CLOCK is due to a decrease in the association rate (k_on_) rather than an increase in the dissociation rate (k_off_) (Figure 3G), consistent with other autoinhibited proteins (25). Furthermore, the k_on_ between full-length CRY1 and CLOCK:BMAL1 PAS-AB (4.73 × 10^4^ M^−1^s^−1^) still falls within the typical diffusion-limited regime (10^4^ – 10^6^ M^−1^s^−1^), rather than the conformational change-limited regime (<10^4^ M^−1^s^−1^) suggesting that autoinhibition of CRY1 by its tail is a short-lived state that occurs within the timescale of diffusion (26).

The transient nature of the CRY1 tail/PHR interaction may also explain the modest change in affinity between full-length CRY1 and constructs of CRY1 that lack exon 11, as the presence of exon 11 increases the equilibrium dissociation constant about 4-fold. While modest, other autoinhibited proteins also demonstrate similar changes in K_*d*_ when their autoinhibitory regions are truncated or removed (27, 28). We showed previously that tuning CRY1 affinity for the BMAL1 TAD within a 4-10-fold range elicited significant period changes in cellular circadian rhythms (3, 4), suggesting that a fold-change in this range is physiologically relevant. To confirm the kinetic analysis, we also measured the equilibrium binding response relative to the concentration of CRY1 titrant (Figure 3H) and found that CRY1Δ11and the PHR have a higher affinity than full-length CRY1 (Figure 3I, J). Therefore, both kinetic and steady-state analysis show that CRY1Δ11 and CRY1 PHR have similar affinities for CLOCK, thus emphasizing the essential role of the tail, and exon 11 in particular, as an autoinhibitory module that regulates CRY1 association with CLOCK:BMAL1.

ChIP-sequencing data of native clock proteins throughout the day in mouse liver revealed that CRY1 participates in two temporally distinct repressive complexes: an ‘early’ complex at circadian time (CT) 16-20, when CRY1 is found in a large complex with CRY2, PER proteins and other epigenetic regulators (29), and a ‘late’ complex at CT0-4 where CRY1 is bound in the absence of other core clock proteins to CLOCK:BMAL1 on DNA (30). Therefore, CRY1 has a distinct role in the clock, in line with studies showing that cellular circadian rhythms require CRY1 (31), and that CRY1 likely works as a repressor both with PER proteins and independently of them (32). PER2 associates with the PHR of CRY1 through its C-terminal CRY-binding domain (CBD) (33). Crystal structures revealed that the CBD of PER2 wraps around the CRY PHR to make contacts near both of the CLOCK and BMAL1 binding sites (Figure 4A) (11, 34). Therefore, we wanted to see if PER2 binding would affect regulation of CRY1 by its tail. First, we examined binding of the fluorescently labeled tail to either CRY1 PHR or a preformed CRY1 PHR:PER2 CBD complex, and found that the presence of PER2 on the PHR decreased tail binding by ~10-fold (Figure 4B). These data suggest that either the PER2 CBD or the tail can bind to CRY1, but not both. We further tested this by titrating the PER2 CBD into a preformed complex of the CRY1 PHR with a fluorescently-labeled tail to show that it displaces the CRY1 tail from the CRY1 PHR (Figure 4C). Therefore, binding of the PER2 CBD and the CRY1 tail to the PHR appear to be mutually exclusive. Given the important role of Exon 11 in maintaining a normal circadian period (7), this suggests that tail regulation of CRY1 function occurs in the late repressive complex at CT0-4 that represses CLOCK:BMAL1 independently of PER proteins (30, 32).

**Fig. 4.:**
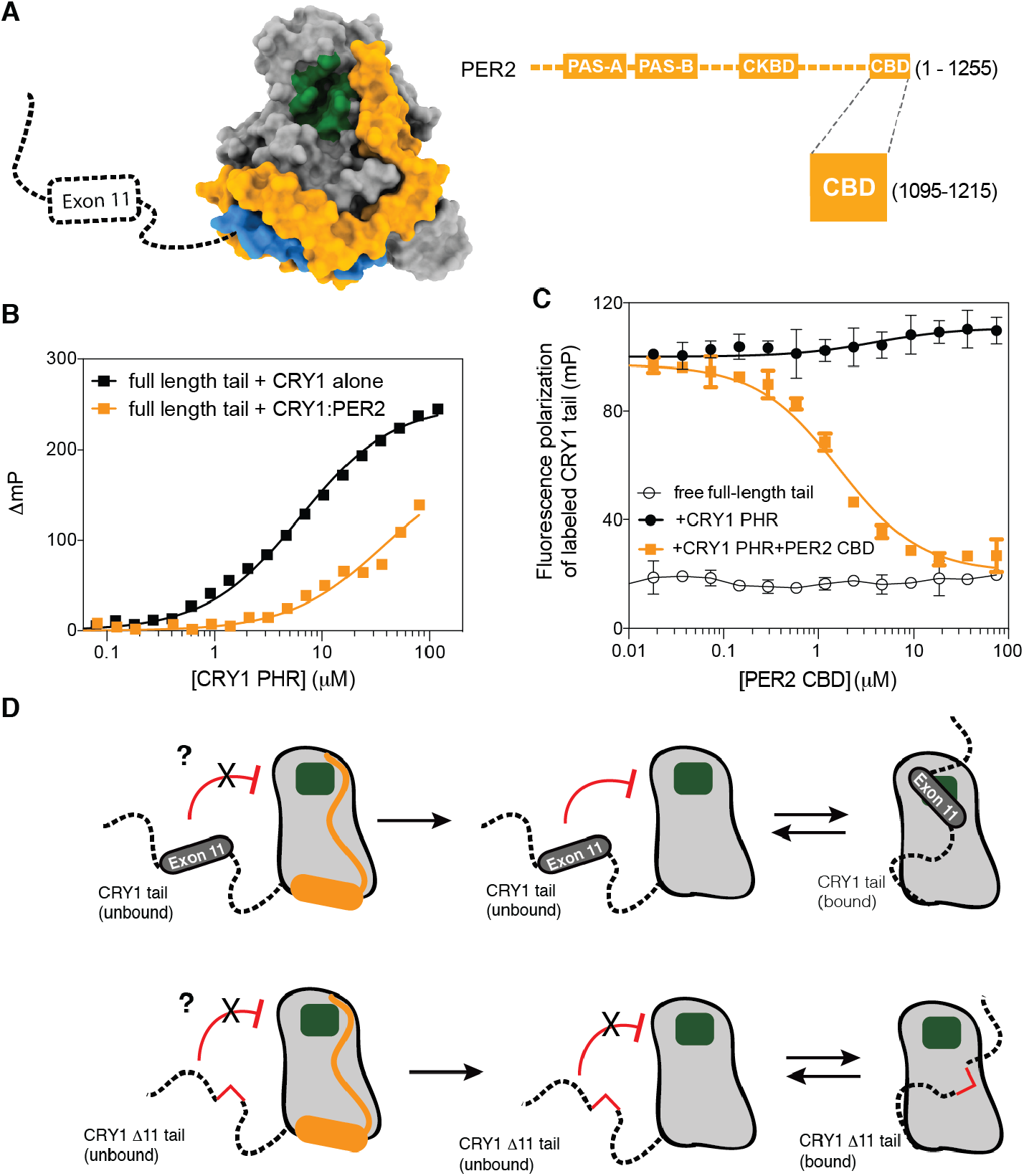
PER2 competes with the CRY1 tail for binding to the PHR domain. (A), Crystal structure of the CRY1 PHR:PER2 CBD complex (PDB: 4CT0) illustrating the proximity of PER2 CBD (orange) to the CLOCK (green) and BMAL1 (blue) binding sites on the CRY1 PHR (gray). (B), FP binding curves of fluorescently-labeled human tail to the CRY1 PHR (black) or a CRY1 PHR:PER2 CBD complex (orange). Plot shows the mean representative binding curves of duplicate samples ± sd (of n = 3 independent assays). Curve represents fit to one-site binding (Prism). (C), FP-based equilibrium competition assay with 4 µM CRY1 PHR bound to 20 nM fluorescently-labeled CRY1 tail in the absence (filled black) or presence of PER2 CBD (orange). Data represent mean ± sd from one representative assay (of n = 3). For assays with displacement, curves represent fit to one-site competitive binding model (Prism). (D), Cartoon model of CRY1 PHR-tail interactions and the role of exon 11 or PER2 in regulating accessibility of the secondary pocket (green).

The data presented here highlight a conserved mechanism between mammalian CRY1 and cryptochromes from *Arabidopsis* and *Drosophila*, where the C-terminal tail reversibly binds to its respective PHR to create an autoinhibited state (Figure 4D and Supplementary Figure 4) (14–16, 20, 35). However, unlike these other cryptochromes that use blue light to regulate the tail/PHR interaction (14, 16, 17, 20), transcriptional repression by mammalian CRY1 does not respond to blue light (36). As an alternative to regulation by phototransduction, we suggest that the association of other proteins (such as PER2), post-translational modifications, and/or regulated alternate splicing of exon 11 could serve to regulate the CRY1 tail/PHR interaction (Supplementary Figure 4). Moreover, exon 11 also shares many attributes with other intrinsically disordered inhibitory modules (idIM) involved in protein autoinhibition; idIMs have a mean length of 24 ± 13 residues (exon 11 is 24 residues) and are often modified in splicing isoforms (CRY1Δ11) (37). The observation that human circadian rhythms are exquisitely sensitive to the absence of CRY1 exon 11 (7) provides support for the growing appreciation that alternate splicing of intrinsically disordered regions is a facile way to modulate protein activity to act as a regulatory mechanism for circadian timekeeping and other cellular signaling systems (38).

## Acknowledgments

We thank Scott Showalter (Penn State) and Isabella Felli (University of Florence) for pulse sequences and advice on ^13^C-detected CON NMR experiments. This work was funded by National Institutes of Health grant R01 GM107069 (to C.L.P.) and funds from the NIH Office of the Director under Award S10OD018455. G.C.G.P. is supported by an HHMI Gilliam Fellowship with additional support from the UCSC Graduate Division.

## Author contributions

G.C.G.P. and C.L.P. designed research; G.C.G.P., I.P., J.L.F., B.N.H., and H.-W.L. performed research; G.C.G.P. and H.-W.L. analyzed data; G.C.G.P. and C.L.P. wrote the paper. All authors approved of the final draft of this manuscript.

## Depositions

The NMR chemical shift assignments for the human CRY1 tail have been deposited in the Biological Magnetic Resonance Bank database with accession number 27988.

## Supplementary Materials

Materials and Methods

Figures S1-S4

References (*39-45*)

