## Supplemental Figures and Methods for "The CRY1 tail controls circadian timing by regulating its association with CLOCK:BMAL1"

This PDF file includes:

- Supplementary Figures S1 to S4
- Materials and Methods
- Supplemental References

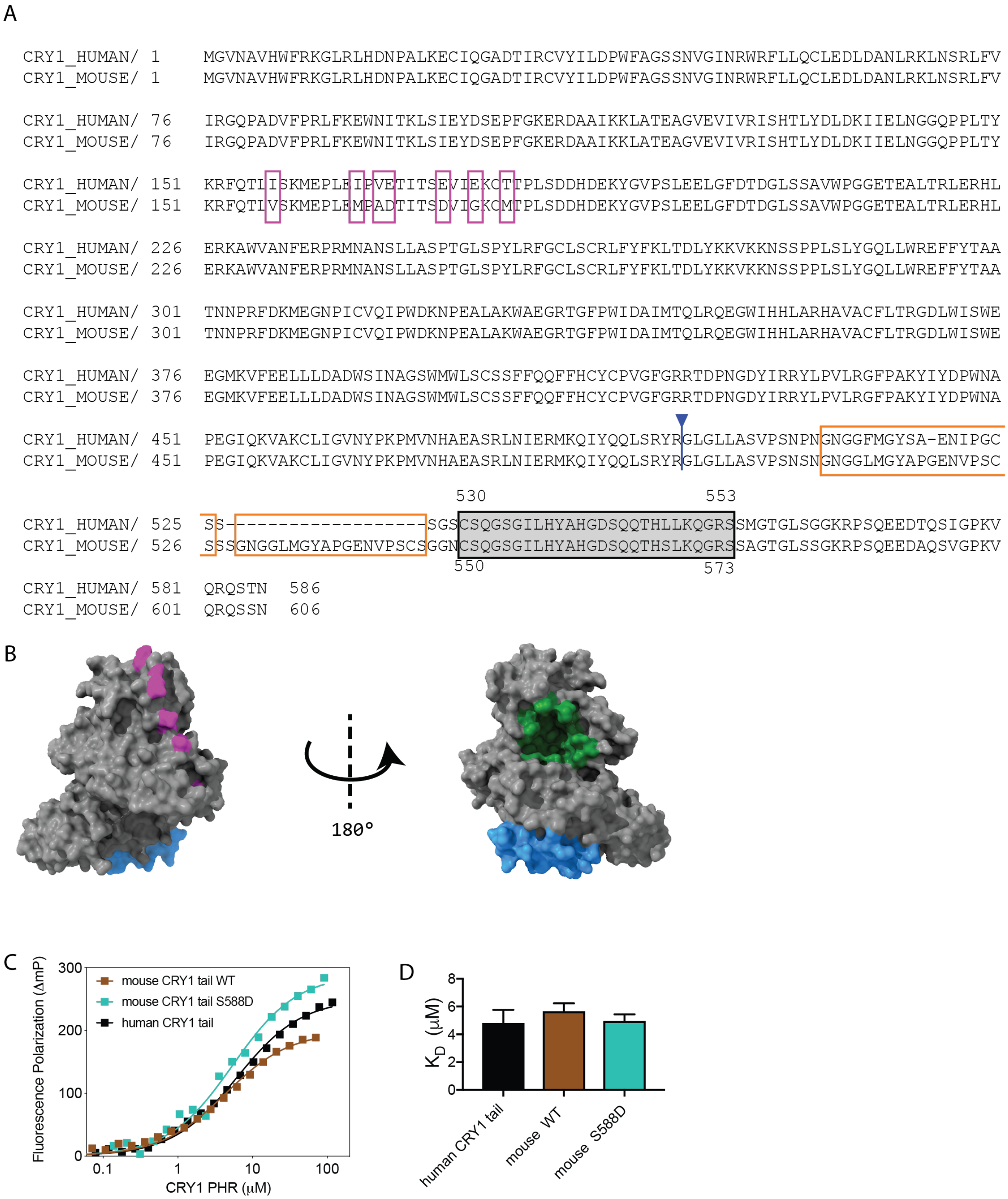


**Fig.S1.**
**Human CRY1 and mouse CRY1 are highly conserved**
A, Sequence alignment of human CRY1 and mouse CRY1. Only 7 residues differ between human and mouse CRY1 PHR (highlighted in purple). The CRY1 tail constructs used in this study start from residue 496 (blue arrow). The mouse CRY1 tail differs from the human CRY1 tail through a repeat insertion at residues at 529 - 546 (highlighted in orange) upstream of exon 11 (gray box, residues 530-553 in human CRY1 and residues 550-573 in mouse CRY1). B, Residues that differ between human and mouse CRY1 PHR are highlighted on the mouse CRY1 PHR crystal structure (PDB ID 5T5X). Non-identical residues (pink) do not overlap with either the CLOCK PAS-B binding site (green) or the BMAL1 TAD binding site (blue). C, FP binding curves of fluorescently-

labeled CRY1 tail constructs to the CRY1 PHR. Plot shows the mean representative binding curves of duplicate samples ± sd (of n = 3 independent assays). Curve represents fit to one-site binding (Prism). D, Affinities of FAM-tail constructs for the PHR derived from FP binding assays (n = 3 independent assays ± sd).

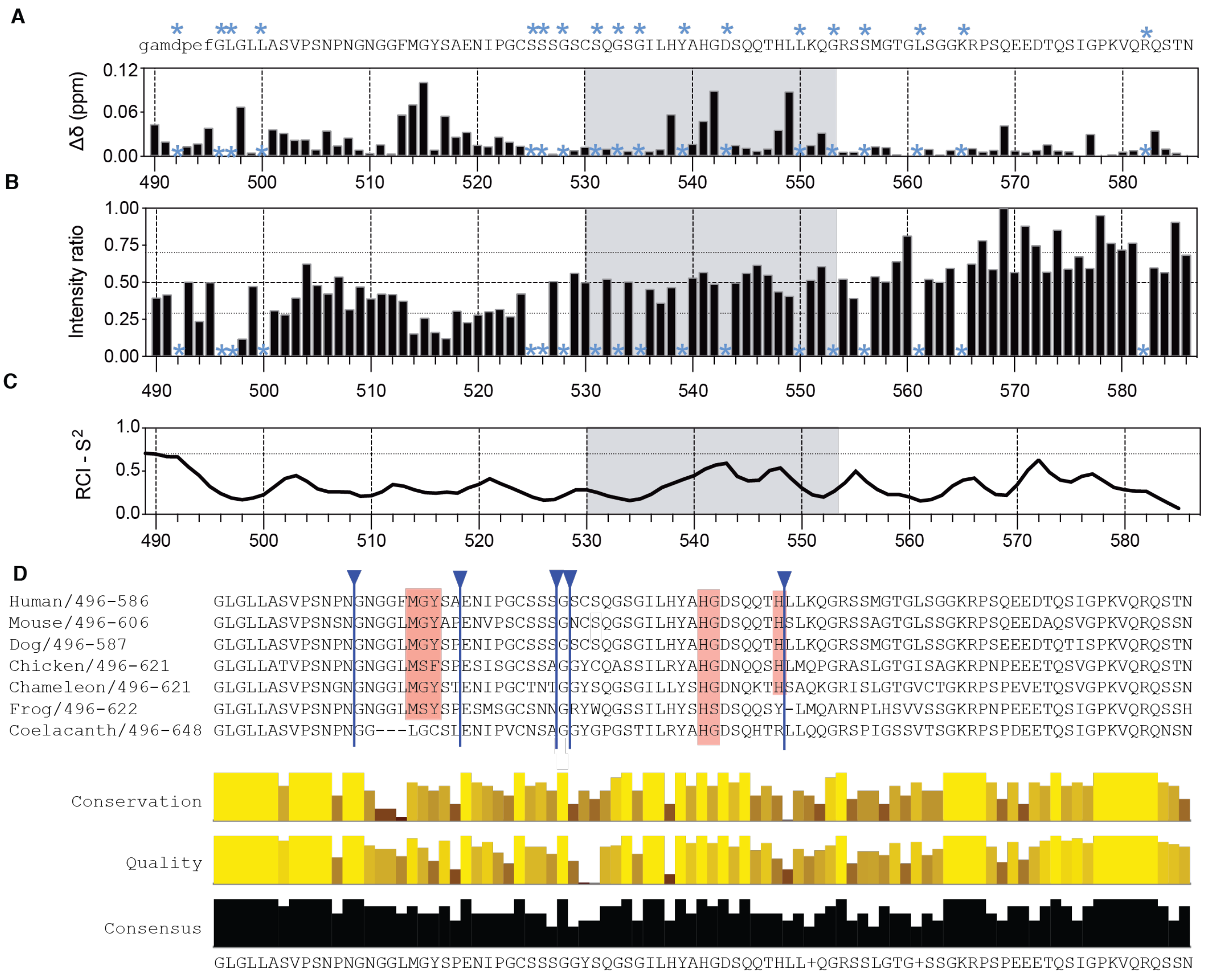


### Fig.S2.

**NMR spectroscopy maps the PHR binding site on the CRY1 tail.**

A-B, Chemical shift perturbation (CSP, A) and relative intensities (B) in the $^{13}$C, $^{15}$N-labeled CRY1 tail including cloning scar (lower case) upon addition of equimolar CRY1 PHR. Asterisk, unassigned or overlapping peaks excluded from the analysis. Exon 11 is represented by a gray box. In H, dashed horizontal line represents the mean intensity ratio ($I_{tail+PHR}/I_{tail}$ alone), while the dotted lines represent sem. Peaks are referred to by the number of the nitrogen of the $^{13}$C-$^{15}$N peptide bond. C, CSI prediction of the lack of secondary structure from NMR data ($S^2 <$ 0.7). D, Sequence alignment and conservation of CRY tail sequences: human (Uniprot ID Q16526); mouse (P97784); Dog (E2RMX4); Chicken (Q8QG61); Chameleon (G1KQX0); Frog (F7B4K1); Coelacanth (H3AHZ5). Potential hotspot residues mapped by NMR are conserved (red box). Insertions that do not align to the human CRY1 tail sequence are hidden and marked by blue arrows: first blue arrow from the left (Chicken residues 509-510; Chameleon residues 509-510; Frog residues 509-510; Coelacanth residue 518); second blue arrow (Mouse residue 519; Dog residue 519; Chicken residue 521; Chameleon residue 521; Frog residue 521; Coelacanth residue 518); third blue arrow (Mouse residues 529-547; Chicken residues 531-500; Chameleon residues 531-550; Frog residues 531-550; Coelacanth residues 528-547); fourth blue arrow (Chicken residues 552-563; Chameleon residues 552-563; Frog residues 552-565; Coelacanth residues 549-588); Fifth blue arrow (Coelacanth residues 609-610). Conservation score calculated by Clustal Omega (39).

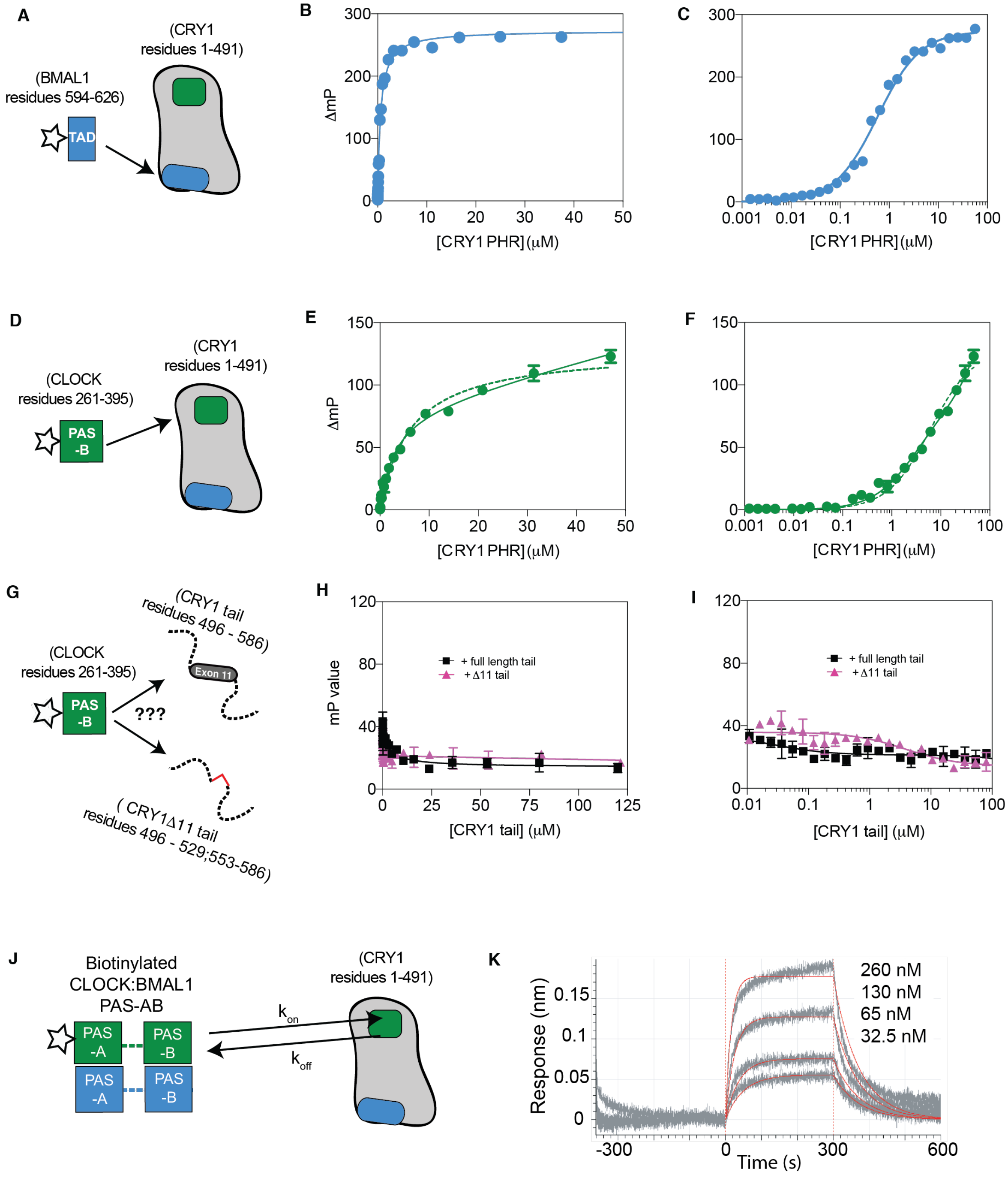


**Fig.S3.**

CRY1 PHR interacts with CLOCK:BMAL1 at distinct sites.

A, The BMAL1 TAD (blue rectangle; fluorescent label, star) binds to the CC-helix (blue cylinder) of the CRY1 PHR (gray). B-C, FP binding curves of fluorescently labeled BMAL1 TAD to the CRY1 PHR in linear scale B and logarithmic scale C. Data fitted to a single site-specific binding model. D, CLOCK PAS-B (green square; fluorescent label, star) binds to the secondary pocket (green rounded square) of the CRY1 PHR (gray). E-F, FP binding curves of fluorescently labeled CLOCK PAS-B to the CRY1 PHR in linear scale E and logarithmic scale F. Data fitted to both single site specific binding model (dashed green line) but some degree of non-specific binding was accounted for by fitting to a single site total model (solid green line). G, CLOCK PAS-B (green square; fluorescent label, star) does not bind to the CRY1 tail (dashed line). H-I, FP binding curves of fluorescently labeled CLOCK PAS-B to the full-length tail (black squares) and Δ11 tail (magenta triangles) in linear scale H and logarithmic scale I depicting insignificant changes in FP upon titration of CRY1 tail. Plots show the mean representative binding curves of duplicate samples ± sd (of n = 3 independent assays) and were fitted on GraphPad Prism. J, CLOCK:BMAL1 PAS-AB (CLOCK, green; BMAL1, blue; biotin, star) binds with CRY1 PHR (gray) through an interaction between CLOCK PAS-B and the secondary pocket (green rounded square). K, BLI sensogram for biotinylated CLOCK:BMAL1 PAS-AB titrated with CRY1 PHR (gray). Model fit represented by a thin red line. Fitted $K_D$ is 65 ± 6 nM with an association rate ($k_{on}$) of 2.01 ± 0.4 x $10^5$ $M^{-1}$ $s^{-1}$ and a dissociation rate ($k_{off}$) of 1.31± 0.1 x $10^{-2}$ $s^{-1}$ .

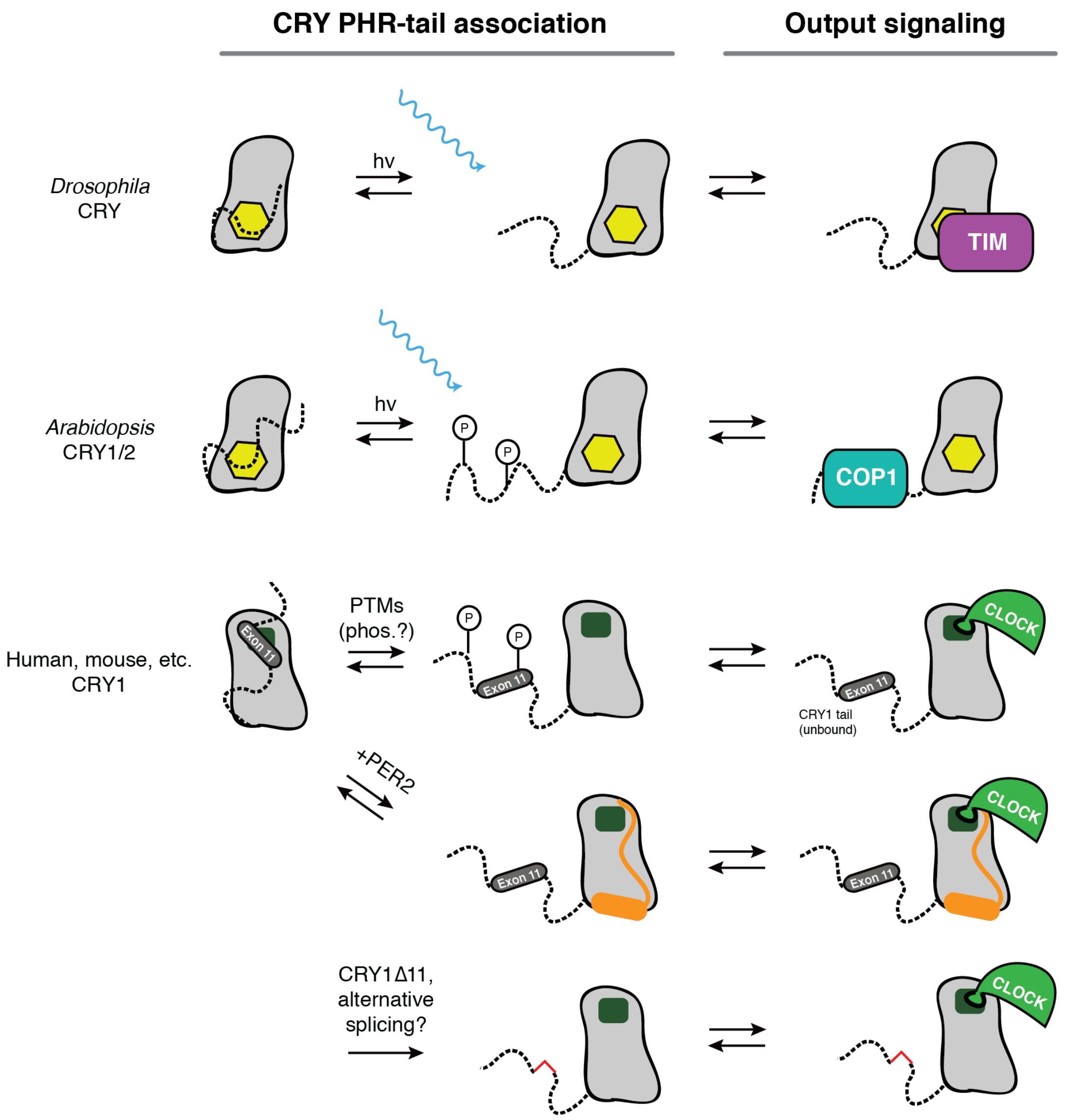


**Fig.S4.**

**The C-terminal of CRY reversibly interacts with its respective PHR to create an autoinhibited state.**

In *Drosophila*, the cryptochrome tail (dashed line) interacts with a flavin within the FAD binding pocket (yellow) of its respective PHR (gray). Blue light induces the release of the tail from its respective PHR thus allowing an interaction between the CRY PHR and circadian regulator Timeless (purple). In *Arabidopsis*, the cryptochrome tail (dashed line) also interacts with a flavin within the cofactor pocket (yellow) of its respective PHR (gray). Blue light induces the release of the tail from its respective PHR thus allowing an interaction between the CRY tail and COP1 (cyan). In mammals, the CRY1 tail possibly interacts with the secondary pocket (dark green) of its respective PHR (gray). Post-translational modifications such as phosphorylation (circled P), or the association of other proteins such as PER2 (orange) could regulate the interaction between the tail and the PHR. When the tail is not bound to the PHR, the secondary pocket (dark green) is open and accessible for binding to the CLOCK PAS-B domain (light green). Gene variants or alternative splicing events that remove exon 11 (red line) have the potential to modulate the tail/PHR interaction.

### Materials and methods

Expression and purification of recombinant proteins:

All CRY1 tail constructs (full length human CRY1 tail residues 496-586; human CRY1 exon 11 residues 530-553; Δ11 tail residues 496-586 Δexon11; human CRY1 exons 10-11 residues 496 – 553; mouse CRY1 tail 496 - 606) and other proteins such as PER2 CBD (human PER2 residues 1095-1215), CLOCK PAS-B (mouse CLOCK residues 261-395), and BAP-tagged CLOCK PAS-AB (mouse CLOCK residues 93-395) were expressed using *Escherichia coli* (*E. coli*) Rosetta2 (DE3) cells. Sortase A and BirA were expressed in BL21 (DE3) *E. coli*. Proteins were expressed as a fusion to the solubilizing tags GST (for BirA), $His_6$-GST (for CRY1 and PER2 constructs), $His_6$-NusA (for CLOCK PAS-B and BAP-tagged CLOCK PAS-AB), or $His_6$ (for Sortase A). Protein expression was induced at 37°C with 0.5 mM isopropyl-β-D-thiogalactopyranoside (IPTG) at an $OD_{600}$ of ~0.8 and grown for an additional 16 hours at 18°C. Cells were centrifuged at 4°C at 3200 x g, reconstituted in 50 mM Tris pH 7.5, 300 mM NaCl, 5% (vol/vol) glycerol and 5 mM β-mercaptoethanol (BME) and lysed using a microfluidizer followed by brief sonication. After clarifying lysate on a centrifuge at 4°C at 140,500 x g for 1 hour, protein was captured using Ni-NTA affinity chromatography (Qiagen) or Glutathione Sepharose 4B resin (GE Life Sciences). Sortase A was prepped in 50 mM Tris, pH 7.5, 150 mM NaCl, 10% (vol/vol) glycerol. GST-BirA was prepped in 50 mM Tris, pH 8.0, 300 mM NaCl, 1 mM dithiothreitol (DTT), 5% (vol/vol) glycerol. For long-term storage, small aliquots of the protein constructs above were immediately frozen in liquid nitrogen and stored at -70°C.

BMAL1 PAS-AB (mouse BMAL1 residues 136-441) and all CRY1 constructs containing the photolyase homology region (mouse CRY1 PHR residues 1-491; full-length mouse CRY1 residues 1-606; full-length human CRY1 residues 1-586; human CRY1Δ11 residues 1-586 Δ530-553) were expressed in Sf9 suspension insect cells (Expression systems) using the baculovirus expression system. $His_6$-mCRY1 PHR, $His_6$-hCRY1 full length, and $His_6$-hCRY1Δ11 were cloned into pFastBac HTa vectors that were later transduced into baculovirus. We used P3 virus to infect Sf9 cells at 1.2 x $10^6$ cells per milliliter and grown for 72 hours at 27°C.

CRY1 expressing cells were centrifuged at 4°C at 3200 x g, resuspended in 50 mM Tris pH 7.5, 300 mM NaCl, 5% (vol/vol) glycerol and 5 mM β-mercaptoethanol (BME) and lysed in low concentrations of detergent (0.01% (vol/vol) Triton X-100), Pierce Protease Inhibitor EDTA-free tablets (1 tablet/50mL, Thermo Scientific), and 1mM phenylmethylsulfonyl fluoride (PMSF) using a microfluidizer followed by brief sonication. After clarifying lysate on a centrifuge at 4°C at 140,500 x g for 1 hour, protein was captured using Ni-NTA affinity chromatography (Qiagen). Protein was further purified using ion exchange chromatography preceding size exclusion chromatography (SEC) into CRY buffer (20 mM HEPES pH 7.5, 125 mM NaCl, 5% (vol/vol) glycerol, and 2 mM Tris(2-carboxyethyl)phosphine (TCEP)). CRY1 protein preps were stored on ice at 4°C. Purified full-length $His_6$-hCRY1 and $His_6$-hCRY1Δ11 were used for experiments within 36 hours of purification, while $His_6$-mCRY1 PHR was used within 7 days.

BMAL1 PAS-AB expressing cells were resuspended in BMAL resuspension buffer (50 mM HEPES buffer pH 7.5, 300 mM NaCl, 5% (vol/vol) glycerol and 5 mM BME). Cells were lysed and clarified as described above. The soluble lysate was bound in batch-mode to Glutathione Sepharose 4B (GE Healthcare), then washed in BMAL1 resuspension buffer and eluted with 50 mM HEPES buffer pH 7.5, 150 mM NaCl, 5% (vol/vol) glycerol and 5 mM β mercaptoethanol, 25 mM reduced glutathione. The protein was desalted into 50 mM HEPES buffer pH 7, 150 mM NaCl, 5% (vol/vol) glycerol and 5 mM β-mercaptoethanol using a HiTrap Desalting column (GE Healthcare) and the GST tag was cleaved with GST-TEV protease overnight at 4°C. The cleaved GST-tag and GST-tagged TEV protease was removed by Glutathione Sepharose 4B (GE Healthcare) and the remaining BMAL1 PAS-AB protein was further purified by Superdex75 gel filtration chromatography (GE Healthcare) into 20 mM HEPES buffer pH 7.5, 125 mM NaCl, 5% (vol/vol) glycerol, and TCEP.

Biotinylation and reconstitution of Biotin-CLOCK:BMAL1 PAS-AB:

For the biotinylation reaction, 100 μM BAP-CLOCK PAS-AB in 20 mM HEPES pH 7.5, 125 mM NaCl, 5% (vol/vol) glycerol, and 2 mM TCEP was incubated at 4˚C overnight with 2 mM ATP, 1 μM GST-BirA and 150 μM biotin. GST-BirA was removed after the reaction using Glutathione Sepharose 4B (GE Healthcare) resin and excess biotin was separated from the labeled protein by gel filtration chromatography. Biotin-CLOCK:BMAL1 PAS-AB heterodimer was reconstituted after labeling by adding equimolar BMAL1 PAS-AB to biotinylated CLOCK PAS-AB and verifying complex formation by SEC.

Fluorescent labeling:

For peptides with shorter sequences such as CRY1 exon 11, BMAL1 TAD, and the Sortase A recognition motif (LPXTG), we utilized synthesized peptides with N-terminal cysteines conjugated to tetramethylrhodamine (TAMRA) 5-maleimide fluorophore. To fluorescently label larger recombinant proteins, we utilized Sortase A-mediated reactions (40) between a fluorescein-labeled (or TAMRA-labeled) LPETGG peptide and our protein of interest (e.g. CRY1 tail or CLOCK PAS-B). CLOCK PAS-B should be labeled right after purification via SEC (i.e. no freeze/thaw cycles). Reactions were carried out in 50 mM Tris, pH 7.5, 150 mM NaCl, and 10 mM $CaCl_2$ using 5 μM $His_6$-Sortase A and 3-5x molar excess of the fluorescently labeled Sortase A motif peptide relative to the protein to be labeled. Labeled protein was purified from the reaction mixture using Ni-NTA affinity chromatography (Qiagen) and/or followed by SEC. Labeled protein was characterized by fluorescent imaging on an SDS-PAGE gel using a Typhoon imager (GE Healthcare). Extent of labeling was measured through spectrophotometry and calculated using the following equation:

$$\%_{labeled} = \frac{A_{dye}}{\varepsilon_{dye} \times \left(\frac{A_{280} - \left(A_{dye} \times CF\right)}{\varepsilon_{protein}}\right)}$$

where $A_{dye}$ is the absorbance at the maximum absorption wavelength (555 nm for TAMRA and 494 nm for fluorescein), $\varepsilon_{dye}$ is the extinction coefficient of the dye (65,000 $M^{-1}$ $cm^{-1}$ for TAMRA and 68,000 $M^{-1}$ $cm^{-1}$ for fluorescein), and the CF is a correction factor that adjusts for the amount of absorbance at 280 nm contributed by the dye (41). We also measured molecular weights of fluorescent probe using a SciEx QTOF mass spectrometer. All probes used were labeled with at least 60% efficiency.

Fluorescence polarization:

All FP assays were performed in 50 mM Bis-Tris Propane pH 7.5, 100 mM NaCl, 0.05% (vol/vol) Tween, 2 mM TCEP. For direct binding assays, varying amounts of CRY1 PHR were mixed with 0.02 µM of a fluorescently labeled CRY1 tail construct. Reactions were incubated for 10 minutes at room temperature. For displacement assays, 0.02 µM fluorescently-labeled CLOCK PAS-B (or 0.02 µM fluorescently-labeled BMAL1 TAD) were incubated with 4 µM CRY1 PHR for 3 hours on ice. Varying amounts of unlabeled CRY1 tail constructs were mixed with this reaction and incubated for 10 minutes at room temperature. Fluorescence polarization measurements were measured on a Perkin Elmer EnVision 2103 Multilabel plate reader with excitation at 485 nm and emission at 535 nm. The equilibrium dissociation constant ($K_D$) and extent of non-specific binding was calculated by fitting millipolarization level (mp) to a one-site total model in GraphPad Prism using averaged mp values from assays with duplicate samples. $IC_{50}$ values were calculated from displacement assays by fitting the mp level to a one-site competitive binding model in GraphPad Prism, with averaged mp values from assays with duplicate samples. Data shown are from one representative experiment (± sd) of three independent assays.

NMR spectroscopy:

All experiments were performed on a Bruker Avance 800 MHz spectrometer equipped with cryogenic probes. Spectra shown for experiments were collected using 20 mM MES buffer pH 6.8, 100 mM NaCl, 4 mM TCEP, 1 mM EDTA, and 10% (vol/vol) $D_2O$ at 298 K. Spectra were processed using nmrPipe and analyzed with Sparky and CCPNMR (41-44). The backbone assignment of the human CRY1 tail was accomplished using standard NH-edited triple-resonance experiments (HN(CA)CO, HNCO, HNCACB, CBCA(CO)NH) and four-dimensional carbon detection methods such as (HACA)N(CA)CON and (HACA)N(CA)NCO (21). We were able to assign 79 non-ambiguous peaks out of 97 possible residues on the CON spectra. Chemical shift perturbation or change in 2D CON peak position was calculated using the following equation:

$$\Delta\delta = \sqrt{\left(N\Delta_{ppm}\alpha\right)^2 + \left(CO\Delta_{ppm}\right)^2}$$

where $\Delta_{ppm}$ is the change in chemical shift and $\alpha = 0.3$ is the normalization between $^{15}N$ and $^{13}C$ chemical shift ranges (45).

Biolayer interferometry:
All BLI experiments were performed using an 8-channel Octet-RED96e (ForteBio). All BLI experiments were performed in BLI buffer (20 mM HEPES pH 7.5, 125 mM NaCl, 5% (vol/vol) glycerol, 2 mM TCEP). For experiments containing full-length CRY1 or CRY1Δ11, we added 0.5mM EDTA to the BLI buffer. For each experiment, we used 8 streptavidin biosensor tips (ForteBio). All experiments began with reference measurements using unloaded streptavidin tips to establish a baseline in BLI buffer, after which non-specific CRY1 association was measured for 5 minutes in wells containing serial dilutions of CRY1 (e.g. 4 μM, 2 μM, 1 μM, 0.5 μM, 0.25 μM, etc.) or a reference sample well containing no CRY, and then dissociation was subsequently measured for 5 minutes in wells containing BLI buffer. After measuring our initial reference, we repeated the same assay with fresh tips that were loaded with 1.5 – 3 μg/mL biotinylated CLOCK:BMAL PAS-AB dimer using BLI buffer that contained no BSA or Tween-20. Data were processed and fitted using Octet software v.7 (ForteBio). Before fitting, all datasets were reference-subtracted, aligned on the y-axis through their respective baselines, aligned for interstep correction through their respective dissociation steps, and finally smoothened using Savitzy-Golay filtering. For each experiment, at least 4 different concentrations were used to fit association and dissociation globally over the full range of the experiment using a 1:1 binding model in Octet software v.7 (ForteBio). Goodness of fit was determined with $\chi^2$ and $R^2$ tests that conform to the manufacturer's guidelines.
